# Integrated Single-Dose Kinome Profiling Data is Predictive of Cancer Cell Line Sensitivity to Kinase Inhibitors

**DOI:** 10.1101/2022.12.06.519165

**Authors:** Chinmaya U. Joisa, Kevin A. Chen, Matthew E. Berginski, Brian T. Golitz, Madison R. Jenner, Silvia G. Herrera Loeza, Jen Jen Yeh, Shawn M. Gomez

## Abstract

Protein kinase activity forms the backbone of cellular information transfer, acting both individually and as part of a broader network, the kinome. Correspondingly, their central role in signaling implicates kinome dysfunction as a common driver of cancer, where numerous kinases have been identified as having a causal or modulating role in cancer development and progression. Driven by their importance, the development of therapies targeting kinases has rapidly grown, with over 70 kinase inhibitors approved for use in the clinic and over double this number currently in clinical trials. Given the growing importance of kinase-targeted therapies, linking the relationship between kinase inhibitor treatment and their effects on downstream cellular phenotype is of clear importance for understanding treatment mechanisms and streamlining compound screening in therapy development. In this work, we combine two large-scale kinome profiling data sets and use them to link inhibitor-kinome interactions with cell line treatment responses (AUC/IC50). We then built computational models on this data set that achieve a high degree of prediction accuracy (R^2^ of 0.7 and RMSE of 0.9), and were able to identify a set of well-characterized and understudied kinases that significantly affect cell responses. Further, we validated these models experimentally by testing predicted effects in breast cancer cell lines, and extended the model scope by performing additional validation in patient-derived pancreatic cancer cell lines. Overall, these results demonstrate that broad quantification of kinome inhibition state is highly predictive of downstream cellular phenotypes.

## Introduction

Computational drug screening has recently emerged as a powerful approach to integrate vast amounts of cancer cell line multi-omics data into predictive models, with the goal of predicting downstream phenotypic responses such as growth and viability. Advancing rapidly with recent developments in machine learning, these methods have the potential to predict outcomes for large drug libraries with minimal experimental cost, reducing the number of drug candidates fed into downstream validation efforts. Most current approaches to the prediction of drug response use baseline cell-line multi-omics data (e.g., mutation status, gene expression, copy number variation, etc.) and Quantitative Structure-Activity Relationships (QSAR) to map drug structure characteristics onto their biological phenotypes, but information describing drug-target interactions, especially at the protein level, remain underutilized because of the unique nature of associated data acquisition methods. For example, the recent DREAM challenge[1] hosted by the National Cancer Institute (NCI) for drug-response predictions saw the winning team utilize high throughput drug screening data along with baseline gene expression features[2] and achieved at most 80% accuracy in predicting cell line responses in a binary fashion.

As one of the foundations of cellular information transfer, protein kinases are enzymes that have also shown promise as therapeutic targets, with initial success being found through the development of Imatinib (Gleevec). Drugs that inhibit kinases (“kinase inhibitors”) are now one of the fastest growing clinical drug classes (74 FDA approved as of 2022), but around 1/3rd of all known kinases still have relatively unknown functions and few chemical tools exist to interrogate and expand this knowledge. To explore the potential of the kinome as a therapeutic target, recent work has focused on profiling the full breadth of targets for kinase inhibitors, especially since many inhibitors have significant off target effects as a result of targeting the conserved ATP-binding pocket. Continued improvements in high-throughput assays such as Kinobead/MS[3], KINOMEscan (© DiscoverX), and KiNativ[4] now enable measurement of a given inhibitor’s interactions across 250-500 kinases, providing a snapshot of its effect on the physiological kinome. We refer to this kinome-wide profiling data as the “kinome inhibition state” of a given inhibitor. This ability to generate drug-target interaction data on a large scale for a compound class is relatively unique, providing a novel means to leverage knowledge of off-target effects for drug response prediction.

The DepMap portal database[5] contains thorough multi-omic characterization of ~1000 cancer cell lines of all types, and cell viability measurements for about ~1500 repurposed compounds, ~250 of which are kinase inhibitors. Using this data, we can connect kinase inhibitor phenotypes of cell viability to their “kinome inhibition states” and build models to predict the cellular responses to treatment with different kinase inhibitors. We have previously shown that these kinome states obtained through the Kinobeads assay for clinical inhibitors are predictive of cancer cell viability, and also validated these predictions experimentally[6]. However, the kinobeads assay is unique and requires dedicated lab personnel to run, restricting its use to relatively few labs. In contrast, the KINOMEscan assay is a popular and easily accessible commercial alternative that assays a panel of ~500 native and mutant kinases recombinantly. Large amounts of KINOMEscan data have been deposited online by various groups[7,8], including data for inhibitors developed against understudied kinases. These altogether account for four times as many inhibitors profiled (~800) when compared to the data available from the kinobeads assay, representing a massive expansion of publicly available inhibitor state data. However, due to the uncharacterized nature of the inhibitors in the large KINOMEscan data set, only a small number of them (~40) have been tested in the DepMap screening database, compared to ~200 inhibitors from the kinobeads set.

In this work, we describe a framework to create an integrated kinome inhibition state data set by combining kinobeads and KINOMEscan data, and then leverage the breadth of this data into predictive models. This combined set contains single-dose inhibitor profiling data for a total of ~800 kinases and kinase interacting proteins, spanning almost 1000 kinase inhibitors that target a diverse section of the overall kinome space. When leveraged within a machine learning framework, and supplemented with baseline gene expression data, we are able to predict the sensitivity of ~450 cancer cell lines in the DepMap screening dataset, with a reasonable R^2^ of ~0.7. Using this model, we were able to generate sensitivity predictions for 1.2 million inhibitor-cell line combinations, many of them targeted towards understudied kinases. We then experimentally validated these predictions in well characterized breast cancer cell lines seen by the model, as well as primary derived pancreatic cancer cell lines. We find reasonable agreement between predicted and observed outcomes in most compounds, seeing an expected drop in performance for understudied compounds and unique patient-derived cell lines. Together, these results show that there is a strong and predictive relationship between the state of the kinome (its “kinotype”) and downstream cellular phenotypes, while further suggesting potential opportunities for leveraging computational models in inhibitor therapy design.

## Results

### Creating an Integrated Set of Kinome Profiling Data Across a Wide Chemical Space

Kinase inhibitors have been profiled using a number of assays, but for this study we have focused on a specific subset of kinase inhibitors that have been assayed using the kinobead/MS-based method [9] or the KINOMEscan^®^ (DiscoverX) method. These methods assess their specific kinase targets as well as the magnitude of inhibition of each kinase in response to different inhibitor concentrations[9]. We combined kinome profiling datasets from Klaeger et al (Kinobeads), LINCS[7] (KINOMEscan), and UNC[10] (KINOMEscan), filtering down to profiles measured only at 1uM. For the small amount of overlap between datasets, the mean inhibition value was taken across drug-kinase combinations. Given that both assays measure the engagement of inhibitors to kinases, most of the proteins that appear in assay results are either known kinases or closely associated proteins. Specifically, kinase inhibitor profiles include measurements on all wild-type and phosphorylated kinases (~500), along with a set of associated proteins (~300). As such, we will refer to this data as “kinase inhibition states”, and the profile of each individual drug as its “kinome inhibition state” (fig 1a).

**Fig 1.**
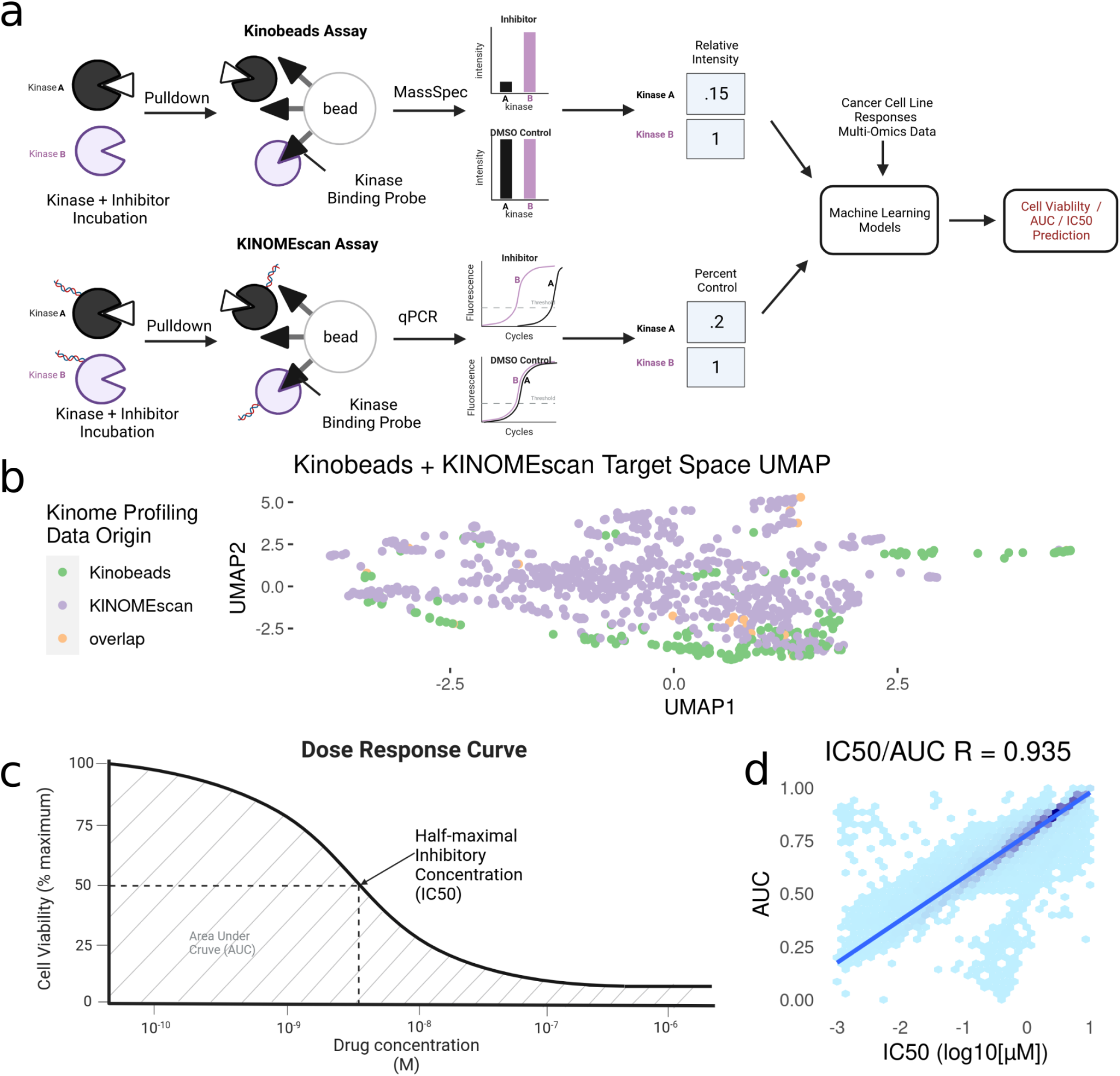
Modeling Pipeline and Target Variable Overview. (a) Schematic of Kinobead/MS (upper) and KINOMEscan assay data integration into machine learning models predicting IC50 and AUC. (b) Visualization of UMAP dimensionality reduction on the combined kinome profiling data set, each point represents a single compound’s position in the target space, and colors representing the origin of kinome profiling data. Target Variables for Modeling: (c) Extraction of IC50 and AUC from a drug’s dose response curve in a given cell line. (d) Correlation and Scales of IC50 vs AUC values. Blue line indicates a linear model fit through the data.

After integration, we were left with a final set of ~1000 compounds with corresponding information on their inhibitor-induced kinome states, describing changes in ~800 kinases and kinase interactors. We then performed a UMAP dimensionality reduction [11] on the dataset for visualization(fig 1b). The UMAP coordinates represent the aggregate effect of each inhibitor on the kinome, i.e. it is a representation of the uniqueness of its kinome inhibition state. Inhibitors that have similar effects on the kinome will have similar coordinates, while disparate inhibitors will have coordinates that are far apart. Using this, we can examine the diversity of kinome space targeting in our dataset, based on the origin of kinome profiling data. Our analysis shows that integrating KINOMEscan data for 800 inhibitors vastly increases the kinome space targeted (fig 1b), compared to just kinobeads alone.

### Connecting Inhibited Kinome States to Cancer Cell Line Sensitivities from the DepMap Repurposing Screen

To connect these kinase inhibitors and their induced inhibition states with their corresponding phenotypes in cancer cell lines, we make use of the DepMap repurposing screen, which uses the PRISM assay[12] to run highly multiplexed cell viability assays. This dataset contains cell viability measurements for over 1500 drugs profiled in 450 cell lines. From within this data, we found ~200 drugs for which we also have corresponding profiling data as described above.

The DepMap repurposing dataset provides cell viability measurements across multiple drug doses, but since our dataset of kinome states is restricted to single-dose measurements, we extracted two single summary statistics for describing cell line sensitivity to kinase inhibitors: Dose-response Area Under the Curve (AUC) and half-maximal Inhibitory Concentration (IC50). These properties are highly correlated with each other, having a Pearson’s correlation coefficient ~ 0.9 (fig 1d). We extracted these properties from DepMap and matched them to our kinome states (fig 1c). The final integrated dataset has ~250 drugs tested across ~450 cell lines, representing ~70,000 inhibitor-cell line combinations representing nearly all cancer types.

### Examining Bivariate Association of Features to Cell Line Sensitivities Provides a Means for Feature Selection

The 450 cell lines tested in the DepMap dataset also have accompanying baseline RNAseq gene expression data, so we integrated the ~20,000 TPM values for each cell line into the kinome-state and cell line sensitivity dataset. This adds baseline cell line-specific gene expression information to our cell line agnostic inhibitor-induced kinome states.

We examined bivariate associations of each of the ~21000 features against the outcome variables (dose response AUC and IC50) using Pearson’s correlation coefficient, and ranked them from largest to smallest absolute value of association. We found that the most correlated feature is the drug-induced kinase inhibition state of TP53RK (fig 2a) with a correlation coefficient R ~0.3, while the most correlated baseline gene expression value was OGFRL1 with a correlation coefficient R ~ 0.05. Overall, inhibitor-induced kinome states showed stronger correlation with cell line sensitivity metrics (Fig 2c) despite there being 40x more baseline gene expression features than kinome states.

**Fig 2.**
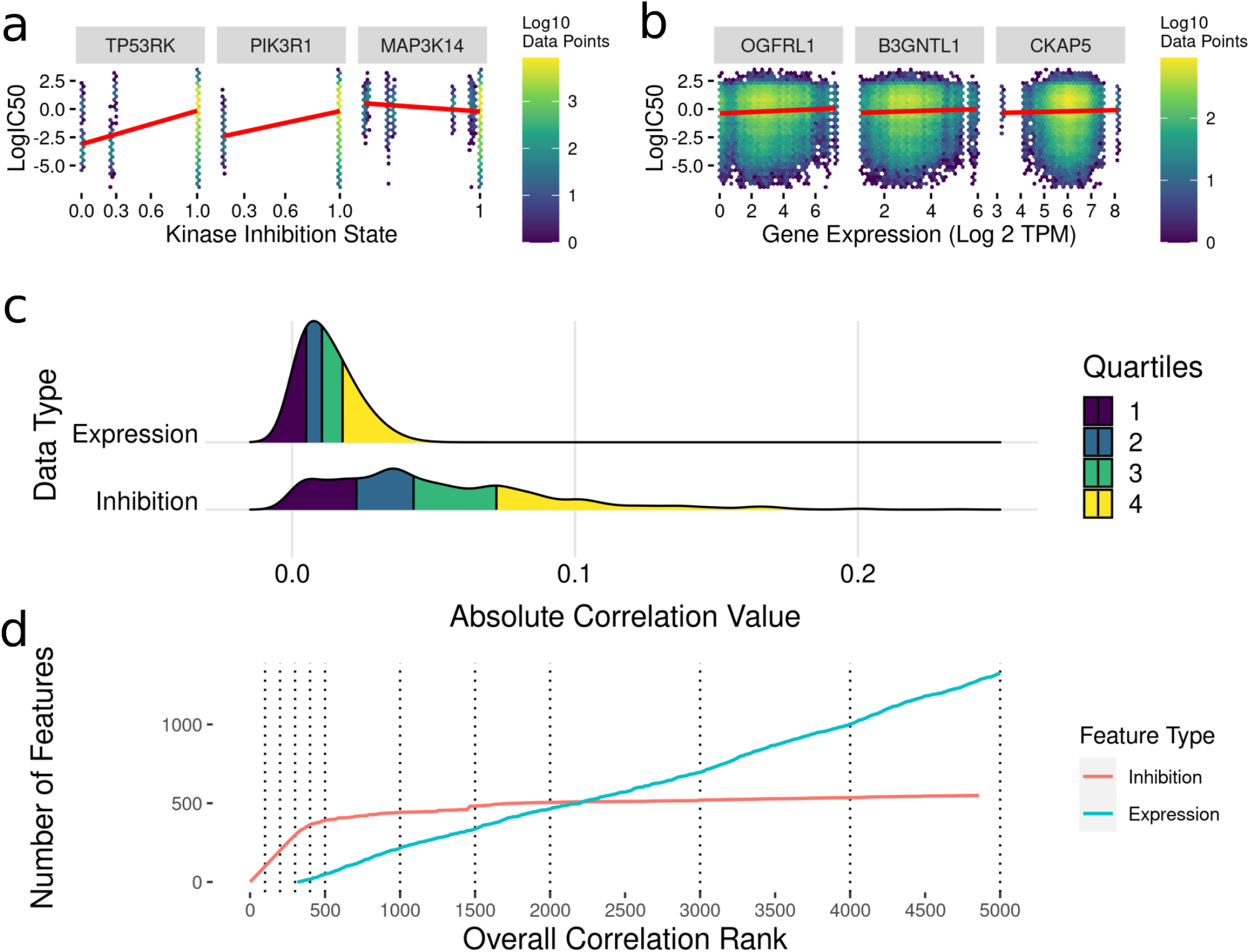
Feature Selection by Bivariate Association with Cancer Cell Line Sensitivity. (a) Sample kinase inhibition state versus LogIC50 heatmap plots showing inhibition states with high (TP53RK), medium (PIK3R1) and low (MAPK14) correlation values. (b) Sample gene expression versus LogIC50 heatmap plots showing genes with high (OGFRL1), medium (B3GNTL1) and low (CKAP5) correlation values. (c) Ridgeline Plot showing distributions of correlations with drug IC50s and AUC values across the data types included in analysis. (d) Plots showing what order classes of features are selected from the ranked set of inhibition states and baseline gene expression values. The dotted lines indicate the discrete increments of feature rank cutoffs at which model performance was tested. Kinase inhibition states were the most informative feature within the first ~300, after which gene expression features started to show predictive value.

After exploring the relationship between each feature and cell line sensitivity, we sought to use machine learning models to combine these features to predict cell line sensitivities to kinase inhibitor treatment. Using the ranked list of feature associations, we utilized a feature selection scheme where we tested discrete increments of the ranked features included in a given model to find the best performers (fig 2d).

### Machine Learning Models can Predict Cancer Cell Line Sensitivity from a Combination of Kinome Inhibition States and Baseline Transcriptomics

To build machine learning models to predict cancer cell line AUC and IC50 in response to treatment with kinase inhibitors, the highest ranked 100-5000 features were selected from the dataset linking drug-induced kinome states to cancer cell line responses (fig 2d).

We compared three model types: LASSO regression, random forest and XGBoost. All models were trained with 10-fold cross validation to minimize overfitting on the training data, ensuring comparable accuracy of the model predictions on new kinase inhibitors and cell lines. For each feature number from 100-5000 we tuned sets of 30 hyperparameters for all model types (fig 3a). The R-squared value between predicted and actual value was utilized as the metric for model comparison. Overall, the 5000 feature XGBoost model performed the best with a cross-validation R-squared of ~0.7 (fig 3b).

**Fig 3.**
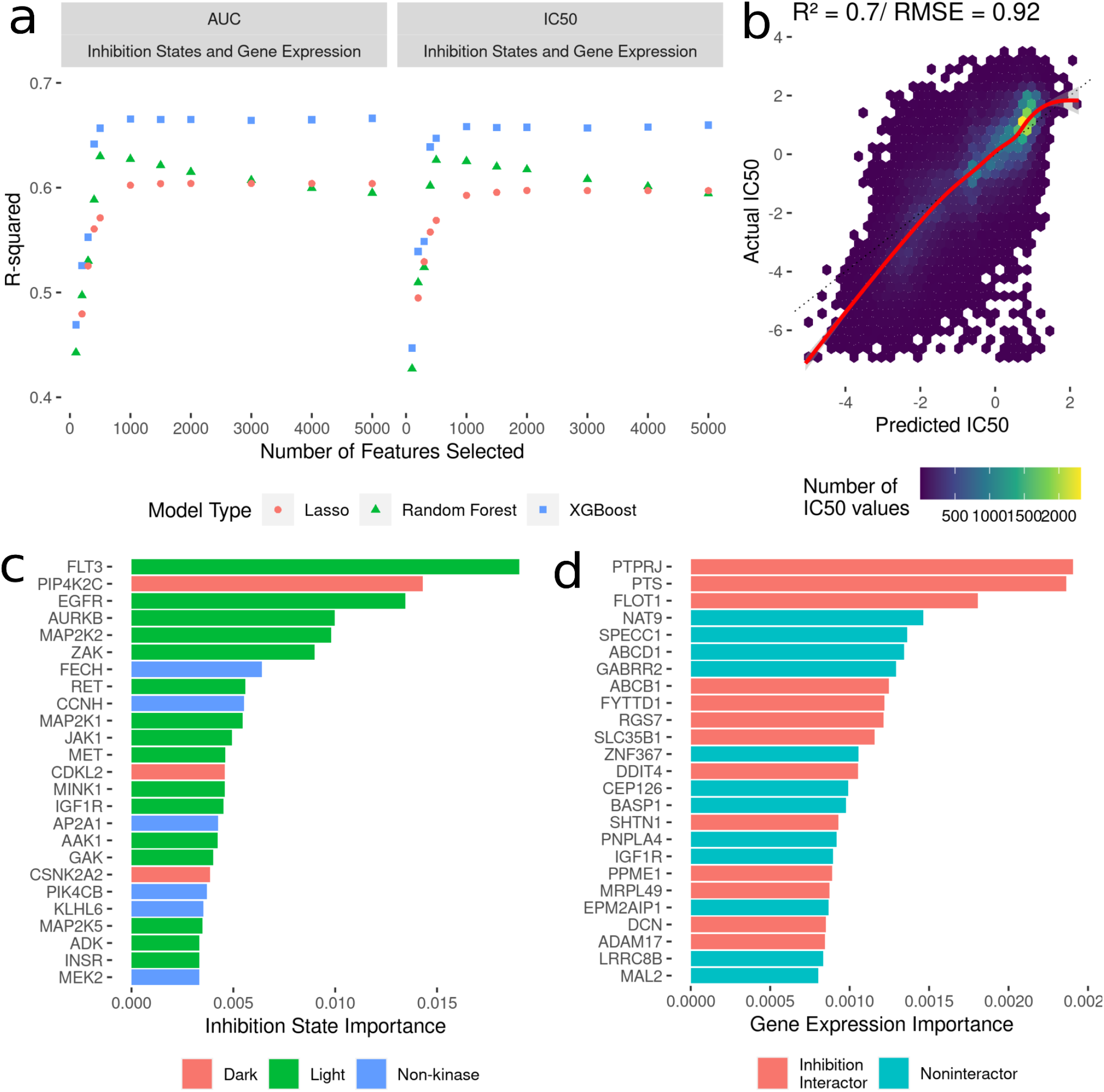
Development of Models to Predict Cancer Cell Line Sensitivities to Kinase Inhibitors by Integrating Single-Dose Kinome Profiling Data. (a) Model performance metrics (R-squared) for LASSO (orange dot), Random Forest (green triangle) and XGBoost (blue square). (b) Scatterplot of predicted IC50 values from the best-performing model vs actual IC50 values. The red line indicated a smooth fit through the data points. (c) Horizontal bar plot showing model importance of individual kinase inhibition states by shapley values. (d) Horizontal bar plot showing model importance of individual baseline gene expression by shapley values.

Since tree-based machine learning models like XGBoost offer in-built explainability, it is possible to interrogate and explain which features were most important in predicting the outcome of cell line sensitivities. These importances generated via shapley values[13] show kinase inhibition states to be overwhelmingly more important for predicting cell line responses when compared to baseline gene expression. Kinases involved in cell cycle and proliferation are overrepresented in the top 25 features (MAP2K, MEK2, CDKL5 etc.), but interestingly six kinase interactor proteins are included as well, suggesting that interactions between inhibitors and non-kinases (off-target effects) have important consequences for cell viability. Baseline gene expression features show much lower model importances, but 40% of the top 25 genes have known interactions with kinases whose inhibition states are used in the model.

### Inclusion of Various Multi-Omics Data with Kinome Inhibition States and Gene Expression Did Not Improve Model Predictive Performance

In addition to the baseline gene expression data, all the cell lines in the DepMap database have three other profiling data types available: copy number variation, gene essentiality from CRISPR/KO, and baseline proteomics. To see if inclusion of these data into models would improve predictions, we integrated these with the modeling dataset of kinome inhibition states and gene expression, and used identical modeling strategies described above to select correlated features, build, and evaluate LASSO, random forest, and XGBoost models (Supp. Fig 1). We found that adding in the various multi-omic data types did not significantly outperform the models limited to kinase inhibition states and baseline gene expression (R-squared of ~0.69 for predicting IC50).

### Experimental Validation of Model Predictions were Successful in Characterized and Novel Cell Lines

After fitting the model on 70,000 cell line-drug combinations, predictions were made on 1.2 million unseen (not seen by the model) drug-cell line combinations. Approximately 90% of the untested inhibitors were associated with KINOMEscan datasets. As an initial validation, we tested a subset of the predictions in well-characterized breast cancer cell lines (HER2 positive: SK-BR-3, BT-474 and two triple negative: SUM159, HCC1806). We analyzed the performance of the model on experimental data for unseen drug-cell line combinations, arriving at an R value ~ 0.6 for all but one (SKBR3) breast cancer cell line(fig 4b). Notably, all the drugs tested had kinome profiling data from the Kinobeads assay.

**Fig 4.**
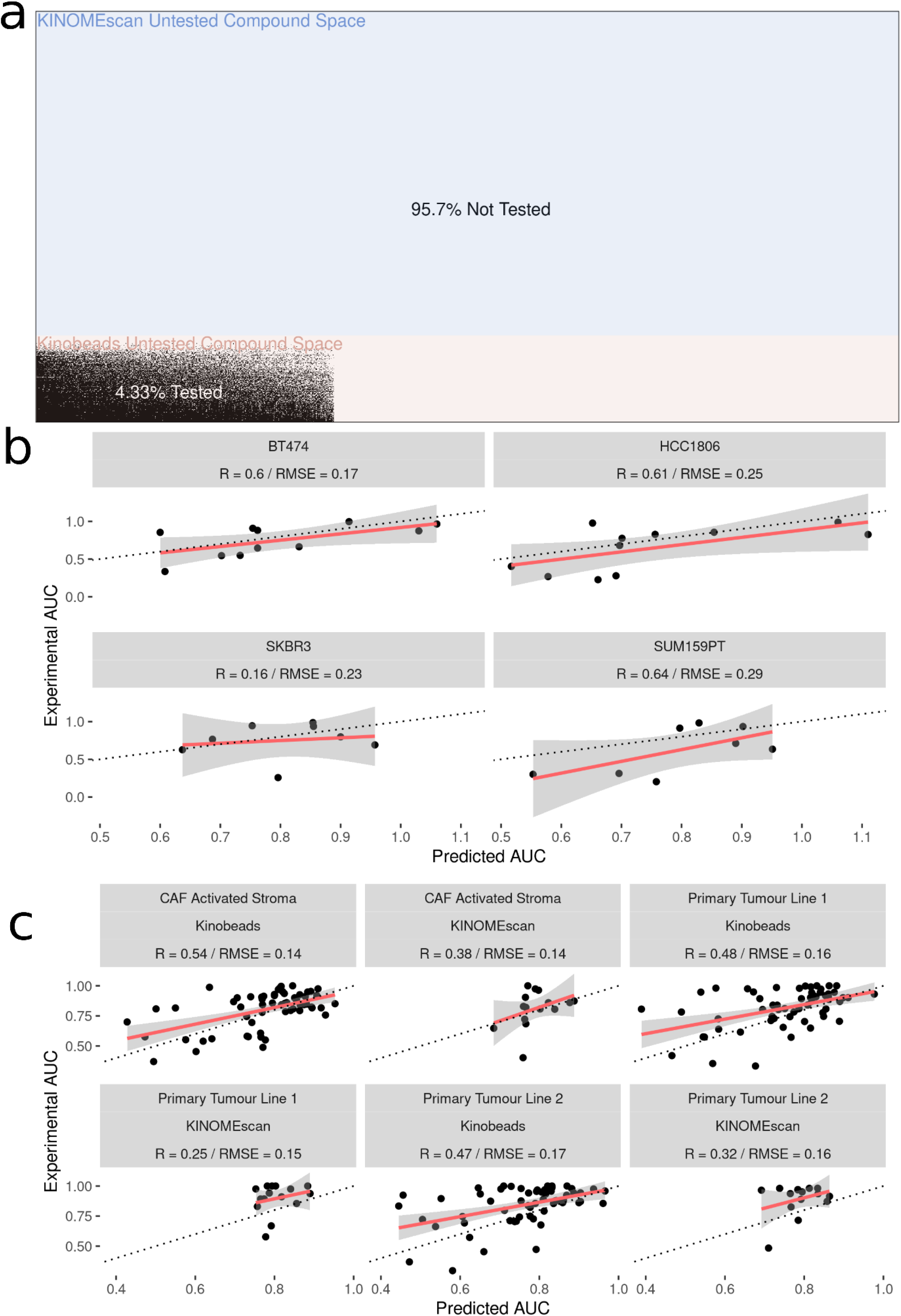
Experimental Validation of Model in Breast Cancer Cell and Patient-Derived PDAC Cell Lines. (a) Visualization of the space of compound (Y-axis) and cell line (X-axis) combinations that have been tested (white) and not tested (black) with colors denoting the origin of the drug kinome profiling data as Kinobeads (Pink) and KINOMEscan (blue). (b) Scatter plot showing relationship between AUC’s predicted by model and experimentally generated AUC’s for drugs not yet tested by PRISM in Breast Cancer Cell Lines (c) Scatter plot showing relationship between AUC’s predicted by model and experimentally generated AUC’s for drugs from Kinobeads and KINOMEscan tested in primary-tumor and stroma PDAC cell lines.

We then further validated the model by predicting inhibitor effects from collected RNAseq data in tumor (two) and stroma (one) derived cell lines from PDAC patients [14,15]. Importantly, these patient-derived cell lines were profiled for baseline gene expression in-house, and represent a novel and highly heterogeneous transcriptional landscape which the model has not seen before. Dose-response AUC predictions were made by the model for 58 drugs with kinome profiling data from the Kinobeads assay and 18 drugs with kinome profiling data from the KINOMEscan assay. The model predicted AUC was compared to experimentally generated AUC, revealing an average R ~ 0.5 for drugs with kinome profiling data from the Kinobeads assay tested in patient stroma-derived cell lines (fig 4c), and R ~ 0.4 for drugs with kinome profiling data from the KINOMEscan assay. On the other hand, in patient tumor-derived cell lines, drugs from kinobeads had model accuracy R ~0.49, and drugs from KINOMEscan had R ~ 0.3 (fig. 4c).

## Discussion

Kinase inhibitors are one of the fastest growing classes of targeted cancer therapies, but only a small fraction of the druggable kinome has been explored to date[16]. We have previously used data describing kinobeads-derived kinome inhibition states to predict cell viability in cancer cell lines in response to clinical kinase inhibitors and shown high prediction accuracy. However, public databases of kinome inhibition states derived through the more accessible KINOMEscan assay cover a large number of uncharacterized compounds and vastly widen the kinome space that can be targeted. In this work, we created a large integrated set of inhibitor-altered kinome states across the kinobeads and KINOMEscan assays, representing a broad space of kinome targets and including a host of tool compounds targeting understudied kinases. We linked these kinome inhibition states to cancer cell line responses to kinase inhibitors (dose response AUC and IC50). We then built machine learning models that integrate these kinome states with cell line baseline gene expression values to predict cell line response to kinase inhibitors. Finally, we predicted cell line sensitivity to previously untested kinase inhibitors in characterized breast cancer and patient-derived PDAC cell lines and validated them experimentally.

Prediction of therapy response for cancer cell lines has been demonstrated through various methods, mostly utilizing chemical structure information, baseline gene expression and gene mutation status. Drug-target interaction data is relatively under-utilized for phenotype prediction, but offers opportunities for biological hypothesis building, especially for compounds with uncharacterized mechanisms of action. Kinome profiling data provides an exciting opportunity to use a wide array of functionally relevant drug-target interactions, and is almost unique (except for GPCR inhibitors and HDAC inhibitors) in terms of ability to assay potential off target interactions. While we have previously shown that kinome profiling data generated from the kinobeads assay is informative for cell viability prediction, KINOMEscan data is much more easily accessible, available publicly for uncharacterized compounds and easier to generate. In addition, it vastly broadens the kinome space capable of being targeted, and increases the number of inhibitors that can be virtually screened as potential therapeutics.

It is important to note that this model linking kinome inhibition states to cell line response is generalizable to any human cancer sample, provided the sample has baseline transcriptomic data available. We have shown in this work and previously that the model can reasonably extrapolate to kinase inhibitors that have not been tested before in well-characterized cell lines. Significantly, in this work we have extended the scope of the model by using novel RNAseq data from patient-derived PDAC cell lines, and testing kinase inhibitors against both tumor and stroma cell lines and achieving reasonable prediction accuracy.

Using tree-based models like gradient boosting lends us the ability to explain to some degree which features most affected cell viability. Shapley importance values generated from the best-performing model show that the inhibition states of kinases had overwhelmingly more predictive power compared to baseline gene expression values, with FLT3 as the most important feature. FLT3 mutations are observed in 30% of Acute Myeloid Leukemia (AML) patients, and various FLT3 inhibitors are commonly prescribed for treatment[17]. However, the study dataset contains a majority of cell lines from Non-small cell lung cancer (NSCLC), and FLT3 inhibitors have recently shown promise in preclinical studies by abrogating DNA damage[18]. Although the baseline gene expression features had considerably lower predictive power in the model, it is important to note that they provided crucial cell line specific context to the model. Interestingly, 40% of all the top 50 gene expression features were annotated as known kinase interactors in the STRING database. An additional strength of modeling cancer response from kinase-drug interactions is the generation of new hypotheses for understudied kinases. For example, the understudied[19] “Dark” kinases PIP4K2C and CSNK2A2 appear in the top 25 features of the best-performing model, suggesting possible functional roles in cancer cell viability. Interestingly, PIP4K2C expression has been associated with outcomes in Acute Myeloid Leukemia (AML)[20], while CSNK2A2 has been associated significantly with prognoses of 14 different cancer types[21].

There are still many limitations to the results reported in this study. The creation of a combined kinome profiling dataset involves gluing together results from different assay types. Although the assays produce the same output (ratio of kinase in treatment sample to kinase in control), there may be numerous methodological artifacts that add noise to the data. We have attempted to address this by analyzing model performance on each assay type individually, and we can see that responses to inhibitors with data originating from KINOMEscan are noisier and more difficult to predict than inhibitors with data from kinobeads. This discrepancy is potentially due to the imbalance in training data availability for the KINOMEscan inhibitors, with only 15 inhibitors having annotated cell line sensitivity data available. In the future, as more cell line sensitivity testing is performed for compounds in the KINOMEscan dataset, model performance for this assay may improve. Additionally, model performance also decreases when shifting from gene expression data from well-characterized cancer cell lines to that of novel patient-derived cell lines. This is potentially due to the innate and significant heterogeneity that exists in such samples and because the models have been trained on baseline transcriptomics set of the given ~450 cell lines.

It should be possible to extend these models to incorporate multiple kinase inhibitors in combination. This is significant, given the frequency of resistance to targeted cancer monotherapies [23] and the potential to escape kinome reprogramming through multi-inhibitor combinations[24]. Thus, another area of future work is to combine the kinome inhibition states of multiple inhibitors to gain an understanding of their dual effect on the kinome, connecting them to biological phenotypes that arise in response to inhibitor combinations to eventually build models and predict effective kinome-targeting combination therapies.

While targeted therapies such as kinase inhibitors have had significant clinical impact, much work remains to better understand how their modulation of the kinome leads to both desirable and undesirable phenotypic effects. The results presented here provide one approach where knowledge of the inhibition state of the kinome can be linked to downstream phenotypes through predictive models, greatly expanding our ability to predict the effects of existing targeted therapies as well as facilitating the design of novel ones.

## Methods

### Data Sources

The primary data sources we used can be split into two categories: the integrated kinome profiling data set and the cancer cell line set:

The following were downloaded from the respective supplementary materials to create the integrated set of kinome profiling data:

1. Kinome profiling data from the kinobeads assay

a. Klaeger et. al 2017
2. Kinome profiling data from KINOMEscan assay

a. LINCS: kinome profiling datasets for individual compounds downloaded programmatically from http://lincs.hms.harvard.edu/db/datasets/ [7]
b. Kinome profiling data for the PKIS drug set was downloaded from the supplementary data of https://journals.plos.org/plosone/article?id=10.1371/journal.pone.0181585 [10]
c. Kinome profiling data for the KCGS drug set was downloaded from the supplementary data of https://www.mdpi.com/1422-0067/22/2/566 [8] and from internal data provided by SGC-UNC.

The following were downloaded from the DepMap portal (https://depmap.org/portal/download/all/) to create the set of cancer cell line sensitivities and their gene expression characteristics:

1. DepMap secondary repurposing screen (“secondary-screen-dose-response-curve-parameters.csv”)
2. CCLE gene expression set (“CCLE_expression.csv”)

The high-throughput screening data gathered in PDAC patient-derived cell lines was gathered from Lipner et al. [14] with methods as described in Berginski et al 2021 [15].

### Data Preprocessing

The scripts implementing these descriptions are all available through github.

#### Klaeger et al. Kinobead Kinase Inhibition Profiles

As previously described [6], we read the values from the supplemental data table into R and produced a filtered list of kinase and kinase interactor relative intensity values. We imputed missing values with the default “no interaction” value of 1, and truncated likely outlier values to the 99.99 percentile (3.43).

#### KINOMEscan Inhibition Profiles

We read in the three datasets mentioned above into R and concatenated them into a single combined data set. All the individual data sets contain identical protein lists because of the same assay type. Values are reported as “Percent Control”, a ratio of protein pulled down in experimental condition (with inhibitor) vs control condition (without inhibitor). These were divided by 100 to convert the scale to 0-1 to match the Kinobeads relative intensity data.

#### Creating the Combined Kinome Inhibition Profiling Set

We took the kinobeads dataset and the KINOMEscan dataset and concatenated them into one large set containing inhibitor-kinase interaction states for ~800 total kinases and kinase interactors. We left out assays that included recombinantly mutated kinases, but left those with naturally occurring post-translational modifications. The vast majority (99.95%) of the inhibitor-kinase pairs represented was unique for either assay type, but for the 0.05% inhibitor-kinase pairs, we took the mean value of the measurements across the two assay types. Additionally, any missing values were imputed with the default “no interaction” value of 1. In the end we were left with kinome inhibition states for ~1000 kinase inhibitors.

#### Dataset of Cancer Cell Line Sensitivity to Drugs from DepMap

The DepMap repurposing dataset contains cell viability measurements across multiple doses, but since our dataset of kinome states is restricted to single-dose measurements, we extracted single summary statistics of cell line sensitivity to kinase inhibitors: Dose-response Area Under the Curve (AUC) and half-maximal Inhibitory Concentration (IC50). We extracted these by reading in the “secondary-screen-dose-response-curve-parameters’’ dataset into R, which contains curve parameters for a log-logisitic curve fit to the cell viability dose response curve and filtered it down to cell line name, IC50, AUC and other associated metadata.

#### Matching of Kinase Inhibitors between Profiling Dataset and Cell-Line Sensitivity Dataset

The compound names from each dataset were read into R, and the package Webchem [25] was used to retrieve PubChem compound IDs. The two sets of compound names were then matched based on these reference IDs. There were 252 matches between the two sets, forming a final set of ~70,000 inhibitor-cell line combinations.

#### Baseline Gene Expression

As described before[6] the RNAseq data provided in the “CCLE_expression.csv” file needed no modifications while preprocessing. Our only modification was to add identifiers to each gene label (“exp_”), to ensure that kinome inhibition data and expression data related to the same gene weren’t accidentally combined.

#### String

The STRING database[26] was processed as described previously[6] to annotate kinases and kinase interacting genes.

### Modeling Techniques

To assess our models we used a random 10-fold cross validation strategy. The number of features was varied as specified by the feature selection scheme described in the results section. We compared the performance of three model types using this strategy: LASSO (Least Absolute Shrinkage and Selection Operator) regression using the glmnet engine[27], random forest using the ranger engine[28] and gradient boosting using the XGBoost (eXtreme Gradient Boosting) engine[29]. Model performance was assessed by the R-squared value between predicted and actual outcome within the cross validation scheme. For each model type, we tuned sets of 30 hyperparameters to find the best possible performer as follows:

1. LASSO

a. Penalty (1E-10 - 0.9)
2. Random Forest

a. Trees (100 - 2000)
3. XGBoost

a. Trees (100 - 1000)
b. Tree Depth (4 - 30)

After final model selection, we fit the model on the entire dataset and then made predictions on inhibitor-cell line pairs not found in the original DepMap screening data.

### Compound Testing

BT-474, HCC1806, SUM-159 and SKBR-3 cells were assayed as described previously [6]. Briefly, cells were grown in ATCC recommended media and seeded at 4000, 2000, 4000 and 500 cells per well respectively. 24 hours after seeding, cells were treated with inhibitors at 30 μM, 3 μM, 1 μM, 300 nM, 100 nM, 30 nM, 10 nM, and 3 nM, along with the appropropriate DMSO controls. seeded at, in white flat-bottom 96-well plates (Corning). Seventy-two hours post-treatment, cells were lysed with CellTiter-Glo (Promega) and luminescence was read using the PHERAstar FS microplate reader (BMG Labtech) and gain adjustments were conducted for each cell line. Data were normalized row-wise to the DMSO-only (0.1% on cells) control samples on each plate to calculate relative viability. Quality checks were performed to look at the data distribution and the presence of spatial bias on a plate. A quality control metric of <120% of DMSO was applied to all rows analyzed.

The functions “ComputeAUC” and “ComputeIC50” from the R package dr4pl [30] was used to fit a four-parameter log-logistic curve to the cell viability data, and extract AUC and IC50 values from the 9-point cell viability curves.

### Software Availability

All of the code written to support this paper is available through github (https://github.com/gomezlab/kinomescan_viability_prediction) along with a walkthrough explaining where to find the code relevant to each part of the paper.

## Acknowledgements

We would like to thank Madison Jenner for providing data access for our PDAC validation. We would like to thank UNC Research Computing for access to the computational resources necessary for this work.

## References

1. Costello JC, Heiser LM, Georgii E, Gönen M, Menden MP, Wang NJ, et al. A community effort to assess and improve drug sensitivity prediction algorithms. Nat Biotechnol. 2014;32: 1202–1212.

2. Gönen M, Margolin AA. Drug susceptibility prediction against a panel of drugs using kernelized Bayesian multitask learning. Bioinformatics. 2014;30: i556–63.

3. Reinecke M, Heinzlmeir S, Wilhelm M, Médard G, Klaeger S, Kuster B. Kinobeads: A chemical proteomic approach for kinase inhibitor selectivity profiling and target discovery. Methods and Principles in Medicinal Chemistry. Wiley; 2019. pp. 97–130. doi:10.1002/9783527818242.ch4

4. Patricelli MP, Nomanbhoy TK, Wu J, Brown H, Zhou D, Zhang J, et al. In situ kinase profiling reveals functionally relevant properties of native kinases. Chem Biol. 2011;18: 699–710.

5. Corsello SM, Nagari RT, Spangler RD, Rossen J, Kocak M, Bryan JG, et al. Discovering the anti-cancer potential of non-oncology drugs by systematic viability profiling. Nat Cancer. 2020;1: 235–248.

6. Berginski ME, Joisa CU, Golitz BT, Gomez SM. Kinome Inhibition States and Multiomics Data Enable Prediction of Cell Viability in Diverse Cancer Types. bioRxiv. 2022. p. 2022.04.08.487646.doi:10.1101/2022.04.08.487646

7. Koleti A, Terryn R, Stathias V, Chung C, Cooper DJ, Turner JP, et al. Data Portal for the Library of Integrated Network-based Cellular Signatures (LINCS) program: integrated access to diverse large-scale cellular perturbation response data. Nucleic Acids Res. 2018;46: D558–D566.

8. Wells CI, Al-Ali H, Andrews DM, Asquith CRM, Axtman AD, Dikic I, et al. The Kinase Chemogenomic Set(KCGS): An Open Science Resource for Kinase Vulnerability Identification. Int J Mol Sci. 2021;22.doi:10.3390/ijms22020566

9. Klaeger S, Heinzlmeir S, Wilhelm M, Polzer H, Vick B, Koenig P-A, et al. The target landscape of clinical kinase drugs. Science. 2017;358. doi:10.1126/science.aan4368

10. Drewry DH, Wells CI, Andrews DM, Angell R, Al-Ali H, Axtman AD, et al. Progress towards a public chemogenomic set for protein kinases and a call for contributions. PLoS One. 2017;12: e0181585.

11. McInnes L, Healy J, Saul N, Großberger L. UMAP: Uniform Manifold Approximation and Projection. J Open Source Softw. 2018;3: 861.

12. Yu C, Mannan AM, Yvone GM, Ross KN, Zhang Y-L, Marton MA, et al. High-throughput identification of genotype-specific cancer vulnerabilities in mixtures of barcoded tumor cell lines. Nat Biotechnol. 2016;34: 419–423.

13. Rozemberczki B, Watson L, Bayer P, Yang H-T, Kiss O, Nilsson S, et al. The Shapley Value in Machine Learning. arXiv [cs.LG]. 2022. Available: http://arxiv.org/abs/2202.05594

14. Lipner MB, Peng XL, Jin C, Xu Y, Gao Y, East MP, et al. Irreversible JNK1-JUN inhibition by JNK-IN-8sensitizes pancreatic cancer to 5-FU/FOLFOX chemotherapy. JCI Insight. 2020;5.doi:10.1172/jci.insight.129905

15. Berginski ME, Jenner MR, Joisa CU, Herrera Loeza SG, Golitz BT, Lipner MB, et al. Kinome state is predictive of cell viability in pancreatic cancer tumor and stroma cell lines. bioRxiv. 2021. p.2021.07.21.451515. doi:10.1101/2021.07.21.451515

16. Laufer S, Bajorath J. New Horizons in Drug Discovery - Understanding and Advancing Different Types of Kinase Inhibitors: Seven Years in Kinase Inhibitor Research with Impressive Achievements and New Future Prospects. J Med Chem. 2022;65: 891–892.

17. Antar AI, Otrock ZK, Jabbour E, Mohty M, Bazarbachi A. FLT3 inhibitors in acute myeloid leukemia: ten frequently asked questions. Leukemia. 2020;34: 682–696.

18. Ryu H, Choi H-K, Kim HJ, Kim A-Y, Song J-Y, Hwang S-G, et al. Antitumor Activity of a Novel Tyrosine Kinase Inhibitor AIU2001 Due to Abrogation of the DNA Damage Repair in Non-Small Cell Lung Cancer Cells. International Journal of Molecular Sciences. 2019. doi:10.3390/ijms20194728

19. Essegian D, Khurana R, Stathias V, Schürer SC. The Clinical Kinase Index: A Method to Prioritize Understudied Kinases as Drug Targets for the Treatment of Cancer. Cell Rep Med. 2020;1: 100128.

20. Lima K, Coelho-Silva JL, Kinker GS, Pereira-Martins DA, Traina F, Fernandes PACM, et al. PIP4K2A and PIP4K2C transcript levels are associated with cytogenetic risk and survival outcomes in acute myeloid leukemia. Cancer Genet. 2019;233-234: 56–66.

21. Strum SW, Gyenis L, Litchfield DW. CSNK2 in cancer: pathophysiology and translational applications. Br J Cancer. 2022;126: 994–1003.

22. Lachmann A, Torre D, Keenan AB, Jagodnik KM, Lee HJ, Wang L, et al. Massive mining of publicly available RNA-seq data from human and mouse. Nat Commun. 2018;9: 1366.

23. Lovly CM, Shaw AT. Molecular pathways: resistance to kinase inhibitors and implications for therapeutic strategies. Clin Cancer Res. 2014;20: 2249–2256.

24. Yesilkanal AE, Johnson GL, Ramos AF, Rosner MR. New strategies for targeting kinase networks in cancer. J Biol Chem. 2021;297: 101128.

25. Szöcs E, Stirling T, Scott ER, Scharmüller A, Schäfer RB. webchem: An R Package to Retrieve Chemical Information from the Web. J Stat Softw. 2020;93: 1–17.

26. Szklarczyk D, Gable AL, Nastou KC, Lyon D, Kirsch R, Pyysalo S, et al. The STRING database in 2021:customizable protein-protein networks, and functional characterization of user-uploaded gene/measurement sets. Nucleic Acids Res. 2021;49: D605–D612.

27. Friedman J, Hastie T, Tibshirani R. Regularization Paths for Generalized Linear Models via Coordinate Descent. J Stat Softw. 2010;33: 1–22.

28. Wright MN, Ziegler A. ranger: A Fast Implementation of Random Forests for High Dimensional Data in C++ and R. J Stat Softw. 2017;77: 1–17.

29. Chen T, Guestrin C. XGBoost: A Scalable Tree Boosting System. arXiv [cs.LG]. 2016. Available:http://arxiv.org/abs/1603.02754

30. Gadagkar SR, Call GB. Computational tools for fitting the Hill equation to dose-response curves. J Pharmacol Toxicol Methods. 2015;71: 68–76.

