## Supplementary Figure 1 for "Integrated Single-Dose Kinome Profiling Data is Predictive of Cancer Cell Line Sensitivity to Kinase Inhibitors"

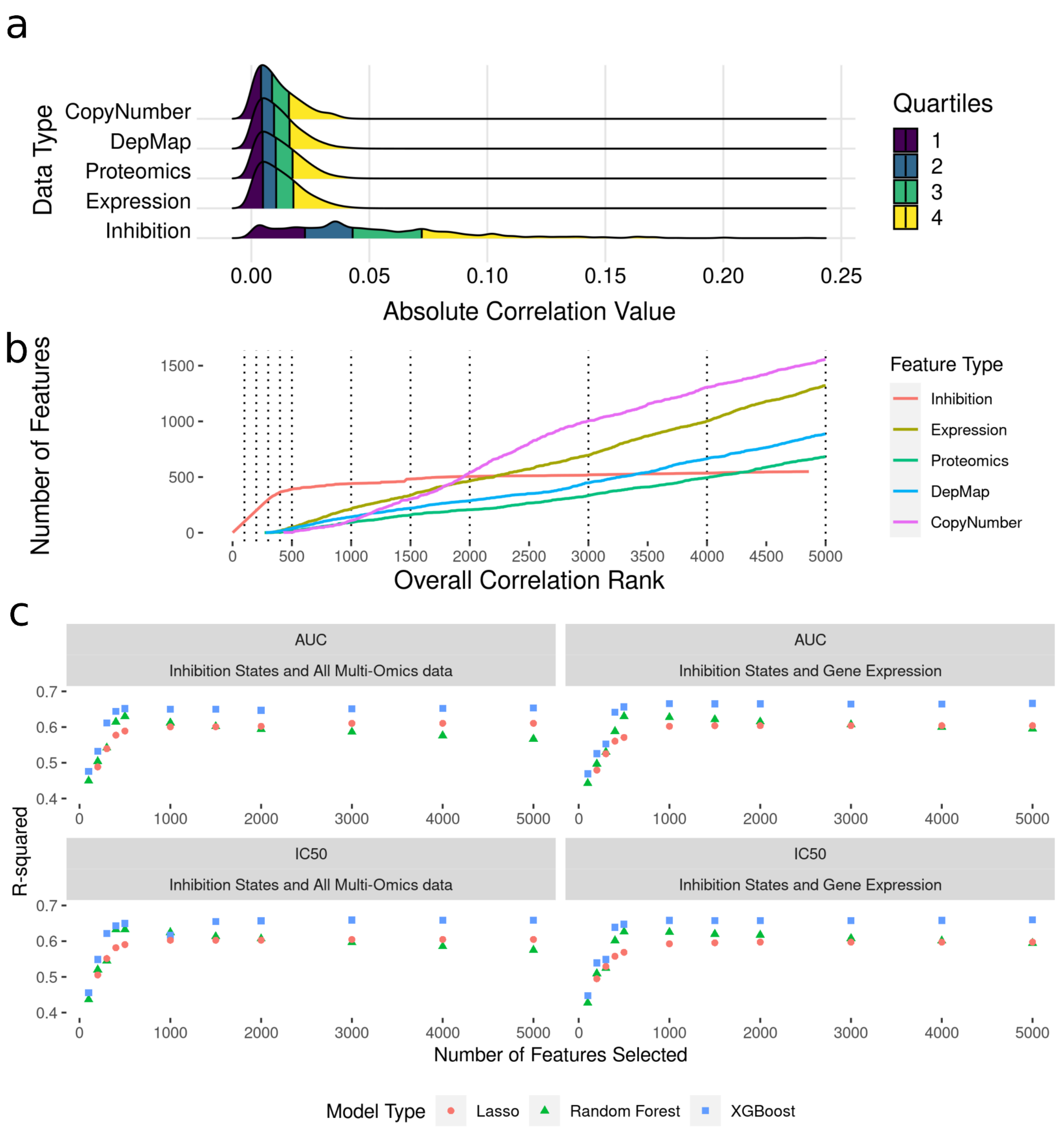


**Supplementary Fig 1. Modeling Results from Integrating Multi-Omics Data with Kinome Inhibition States and Baseline Gene Expression to Predict Cell Line Sensitivity.** (a) Ridgeline Plot showing distributions of correlations with drug IC50s and AUC values across the data types included in analysis. (b) Plots showing what order classes of features are selected from the ranked set of inhibition states and baseline gene expression values. The dotted lines indicate the discrete increments of feature rank cutoffs at which model performance was tested. (c) Model performance metrics (R-squared) for LASSO (orange dot), Random Forest (green triangle) and XGBoost (blue square).
